# Methods for tagging an ectoparasite, the salmon louse *Lepeophtheirus salmonis*

**DOI:** 10.1101/2023.08.31.555695

**Authors:** Alexius Folk, Adèle Mennerat

## Abstract

Monitoring individuals within populations is a cornerstone in evolutionary ecology, yet individual tracking of invertebrates and particularly parasitic organisms remains rare. To address this gap, we describe here a method for attaching radio frequency identification (RFID) tags to individual adult females of a marine ectoparasite, the salmon louse *Lepeophtheirus salmonis*. Comparing two alternative types of glue, we found that one of them (2-octyl cyanoacrylate, *2oc*) gave a significantly higher tag retention rate than the other (ethyl 2-cyanoacrylate, *e2c*). This glue comparison test also resulted in a higher loss rate of adult ectoparasites from the population where tagging was done using *2oc*, but this included males not tagged and thus could also suggest a mere tank effect. Corroborating this, a more extensive analysis using data collected over two years showed no significant difference in mortality after repeated exposure to the *2oc* glue, nor did it show any significant effect of the tagging procedure on the reproduction of female salmon lice. The proportion of RFID-tagged individuals followed a negative exponential decline, with tag retention among the living female population generally high. The projected retention was found to be about 88% after 30 days or 80% after 60 days, although one of the four batches of glue used, purchased from a different supplier, appeared to give significantly lower tag retention and with greater initial loss (74% and 60% respectively). Overall, we find that RFID tagging is a simple and effective technology that enables documenting individual life histories for invertebrates of a suitable size, including marine and parasitic species, and that it can be used over long periods of study.

## Introduction

In the current biodiversity crisis, it is particularly important to understand how interacting species and in particular disease organisms respond to anthropogenic change (Hudson et al., 2006, Smith et al., 2009, Gupta et al., 2020). Disease ecology has developed as a major field over the past decades, revealing the scope and complexity of parasitism’s effects in natural populations (Dobson et al., 2008, Poulin 2014). Comparatively little focus has been given to the selection that hosts and host demography exert on their parasites (Hudson et al., 2002, Mennerat et al., 2010, Ebert & Fields, 2020 – but see e.g. Kennedy et al., 2016, Ugelvik et al., 2017). This is in part due to the study of parasite behaviour and evolution being limited to population-level data, as tracking individual parasites is notoriously difficult.

In non-parasitic species, tracking individuals across life stages has allowed researchers to study the causes and consequences of phenotypic variation within populations (Clutton-Brock & Sheldon 2010). Monitoring populations of uniquely marked individuals is a cornerstone of evolutionary ecology, as accessing individual life histories allows insight into microevolution and dispersal, as well as informing conservation decisions (Clutton-Brock & Sheldon 2010, Bonte et al., 2012, Charmantier et al., 2014, Pemberton et al., 2022, Sheldon et al., 2022).

Studies of individually marked vertebrates, especially birds and mammals, have traditionally been far more common than studies involving invertebrates (Sheldon et al., 2022) due to the difficulty in consistently identifying small, numerous, and mobile animals (Hagler & Jackson 2001, Streit et al., 2003, Rodríguez-Muñoz et al., 2019). A few notable exceptions include long-term longitudinal studies in natural populations of insects (e.g. the field cricket, Rodríguez-Muñoz et al., 2019) and research on lepidopteran dispersal between habitat fragments (e.g. Schtickzelle & Baguette 2003). Recent advances in radio-frequency identification (RFID) technology now also allow for the tagging and digital identification of more invertebrate species, including eusocial insects such as bees and ants (Streit et al., 2003, Robinson et al., 2014, de Souza et al., 2018, Nunes-Silva et al., 2019).

While individual tagging approaches will likely remain difficult for most internal parasites, ectoparasites living on the outer surfaces of their hosts are more easily accessed and thus have a greater potential for individual tracking using similar techniques to those currently used on free-living invertebrates, with similar limitations (Rataud et al., 2020).

Here we describe a method for tracking individual adults of an ectoparasite, the copepod *Lepeophtheirus salmonis* (salmon louse), using an RFID tagging system. *L. salmonis* is a marine species that cannot complete its life cycle in brackish or freshwater. It became a major pest following the exponential growth of Atlantic salmon (*Salmo salar*) populations in marine intensive aquaculture facilities, where it experiences anthropogenic selection (Mennerat et al., 2010, Coates et al., 2021) and contributes to the decline of wild salmonid populations (Costello 2009, Vollset et al., 2016, Shephard & Gargan 2021). In this article, we detail our method for attaching RFID tags onto adult female *L. salmonis*, document the retention time of the tags, and test their effects on mortality and reproduction.

## Methods

### Salmon hosts and culturing of lice

To compare types of glue, document retention rates, and test the effect of RFID-tagging on *L. salmonis* mortality and reproduction, we infected naïve Atlantic salmon (Industrilaboratoriet, Bergen, Norway) using salmon louse larvae hatched from eggs sampled from a marine facility located at the Institute of Marine Research, Austevoll, Norway (glue comparison test), and from commercial salmon farms in Fosså, Rogaland county, Norway and Oppedal, Vestland county, Norway (glue retention time and impacts on mortality and reproduction). Salmon lice eggs were incubated in the lab for 14 days following protocol described in Hamre et al., (2009). The copepodites from those eggs were used to infect post-smolt Atlantic salmon housed in 1m x 1m, 500L tanks supplied with filtered and UV-treated seawater (flowrate, 2 - 6 L min^-1^, temperature, 7.1 - 9.5 °C). The salmon hosts used in the initial glue comparison test were kept in two tanks containing 15 to 16 fish each, while those used in the other tests were kept in three pairs of tanks housing either 15 or five fish. Tanks with either 15 or 16 fish were infected by lowering the water level and adding 30 copepodites per fish on average (following e.g. Mennerat et al., 2017), while tanks with five fish were infected at full tank volume with an average of 15 copepodites per fish, as part of another, ongoing study. The tanks were equipped with sieves to filter outlet water and daily collect salmon lice that had detached from their hosts and been flushed out.

### Handling and registration of salmon lice

From 60 days post-infection onwards, salmon hosts were individually netted and anesthetised until unresponsive in 1 mL L^-1^ metomidate and 0.3 mL L^-1^ benzocaine (glue comparison study), or later on with either these anesthetic compounds or with 120 mg L^-1^ tricaine mesylate (glue retention time and impacts on mortality and reproduction), after which they were inspected for salmon lice. All adult lice were carefully removed with fine curved forceps and placed onto a moistened paper label in a petri dish, after which the salmon was placed into a holding tank to recover. Female salmon lice were tagged with p-Chips using surgical glue (see below for details). Finally, tagged lice were registered by scanning their RFID tags, photographed, and placed back onto their original host, which was returned to its tank. Subsequent checks for lice followed a similar host capture and anesthesia procedure.

### Attachment of RFID tags

For an ecdysozoan like *L. salmonis*, attaching RFID tags externally can only be done on adults, i.e., once they stop moulting. Adult female *L. salmonis* can be tagged by gluing an RFID tag onto the dorsal side of their abdomen (genital segment). To do so we used a ‘p-Chip’ microtransponder (p-Chip Corp., Princeton, NJ, U.S.A.), a 500 x 500 x 100*µ*m RFID tag carrying a unique 9-digit identification number. The readable side of the tag contains a photocell that is powered by a laser on a tag-reading wand, which records the ID number as well as additional information as a comment (e.g. sample name), and logs all scans in a .csv file. In males, which are the smaller sex, the genital segment is too small for a p-Chip, and gluing the tag to the cephalothorax was found to be invariably and quickly fatal.

To prepare female salmon lice for tagging, we set all lice onto a paper label, wetted with cold seawater in a petri dish. The top of the genital segment was thoroughly dried with a cotton swab (**Figure 1a**), and a small amount of glue was applied to the center of the segment by touching it with the tip of a wooden toothpick (**Figure 1b**). The tag was placed facing upward onto the louse’s genital segment using the tip of a clean, damp toothpick (**Figure 1c**), and an additional dot of glue was applied over the edges of the tag to encapsulate it (**Figure 1d**).

**Figure 1.**
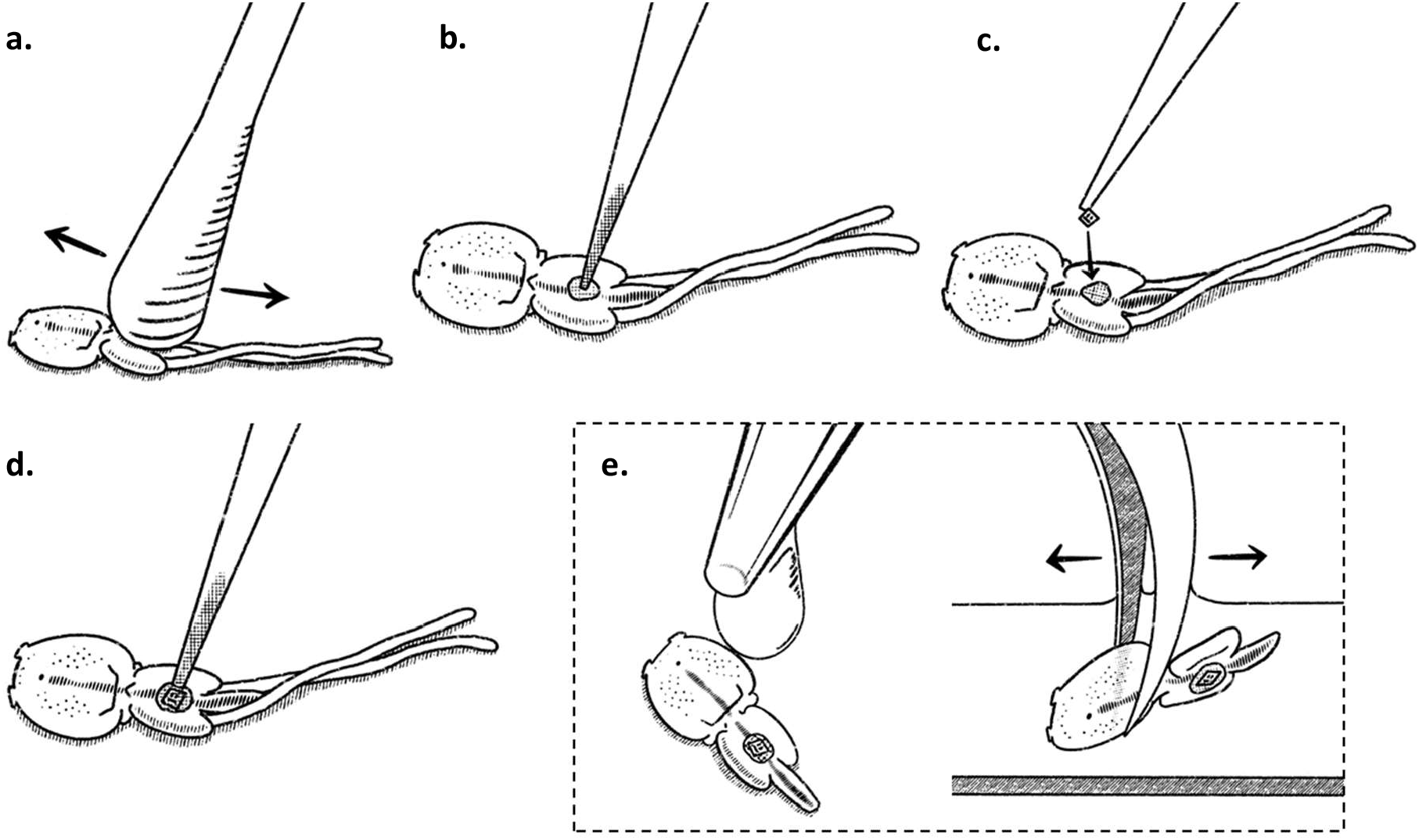
Procedure for tagging female salmon lice with RFID tags. (a) Drying the genital segment with a cotton swab; (b) applying glue to the louse with a toothpick; (c) placing a p-Chip onto the louse with a damp toothpick; (d) applying glue over the top of the p-Chip; and then (e) dripping saltwater onto the louse and/or dipping it in saltwater to set the superglue. (Illustration A. Folk)

To set the glue and following the removal of any extruded egg matrices, cold seawater was either dripped onto the salmon louse, or the louse was gently dipped into the water, or both (**Figure 1e**). Any excess glue that might have detached upon contact with water was removed to prevent clumping around the female’s gonopores. After letting the glue set for an additional 30 - 60 seconds, newly tagged female lice were squirted again with saltwater and then flipped onto their backs to be re-applied to their original host. With some experience, the RFID application process takes two to five minutes for one to nine lice aligned in the same Petri dish. Multiple lice may be prepared in this way simultaneously.

Due to the short transmission distance, p-Chips attached to salmon lice can only be read near the reading wand (< 1cm). Upon later checks of previously tagged lice, we removed them from their hosts, set them onto a wet paper label in a Petri dish, and brought them to the chip reader.

## Video demonstration

https://paranthrope.w.uib.no/files/2023/12/licechip_procedure_v2.mp4

### Comparison of two alternative types of glue

According to a preliminary search, two alternatives for glue seemed to be 2-octyl cyanoacrylate (Surgi-Lock, hereafter 2oc) and ethyl 2-cyanoacrylate (Würth Klebfix, hereafter e2c). Pilot studies did not reveal higher mortality between RFID-tagged and untagged lice glued using either type of glue (see **Table SM1, Figure SM3**). To confirm this and determine the more effective glue, adult female salmon lice in the 15-fish tank (*n* = 62) were tagged with p-Chips using 2oc at 59 days post-infection, while those in the 16-fish tank (*n* = 79) were tagged using e2c at 66 days post-infection.

After the initial RFID-tagging and until the end of the experiment, the tank outlet filters were checked daily for dislodged and dead salmon lice. All fish were re-captured 21 days after initial tagging; salmon lice were sampled and registered following the procedure outlined above and placed back on their hosts. The fish were caught, the lice tallied, and their p-Chips scanned a final time 52 and 59 days post-tagging for the e2c group and the 2oc group, respectively. All lice at every step were recorded as being either male or female, and for the females, as being either tagged or untagged. Since it was possible to capture all female salmon lice during the initial RFID-tagging and no new individuals were added after that, all untagged recaptured females were known to have been tagged previously. At the conclusion of the study, the salmon were humanely euthanised under full anesthesia. One salmon in the e2c group was instead euthanised 6 days prior to the end of the study due to a damaged fin.

Combining the daily checks of the outlet filters and the two recapture sessions, we obtained the numbers of tagged and untagged females for successive days of the study. Between census events, the proportions of females in each tank retaining their RFID tags were estimated using observations from tank outlet filters. Females found in filters were included in the tally of tagged or untagged females for each day, and subtracted from population size for subsequent observations. Tag retention was compared between the two glue types using a generalised linear model (GLM), where the response variable was the log-transformed proportion of retained RFID tags weighted by simultaneous population size. The explanatory variables used were glue type as a factor and time elapsed (in days) since initial tagging as a covariate, as well as the interaction between the two.

The proportions of female and male salmon lice remaining alive were estimated using the number of lice observed during a population census, from which we substracted the sum of lice found in the tank outlet filters over successive days in between census events. The starting population size for each group was the number of adult male and female lice observed on the day of initial tagging (minus three females and two males from a fish euthanized in the e2c tank). Mortality was compared between the two glue types (e2c: 56 untagged adult males, 79 tagged adult females; 2oc: 63 untagged adult males, 62 tagged adult females) using a GLM of the log-transformed proportion as the response variable weighted by starting population size. Explanatory variables included glue type and sex as factors, the number of days since tagging as a covariate, as well as the glue type *\** time and sex *\** time interactions.

### Documenting retention time

Following our glue comparison test, we tagged hundreds of female salmon lice as part of other ongoing research. Our methods for tagging follow the procedure described above using 2-octyl cyanoacrylate (2oc), and the salmon lice are surveyed every six to 10 days. Here we use data collected during a period ranging from 11th January 2021 to 16th December 2022.

All female salmon lice (*n* = 948) were tagged at adulthood, usually after extruding their first clutch (59.2 ± 0.15 SE days post-infection). On subsequent checks, adult females found without a readable p-Chip were tagged again and their IDs were determined by visually comparing them to pictures taken previously, as their pigmentation patterns are repeatable and unique (see **Figure SM1**). The time until tag loss is the number of days between the initial tagging date and the date of the first check when the p-Chip was observed to be missing (212 instances) or nonfunctional (4 instances). For those lice that were re-tagged multiple times, only the retention of original p-Chips is considered here. It was difficult to positively identify all dead and detached lice in the outlet filter without p-Chips as some of those found were too damaged to be identified from pictures. As a result, the date of mortality was determined either by the date at which we found a tagged louse in the tank’s outlet filter, or else as the date of the following weekly check when the louse was no longer observed.

Starting from day 0 (the date of tagging), the data were split into 7-day periods in which all adult females alive at that time were tallied as having either retained or lost their original p-Chip (overview in **Figure SM2**). To estimate the daily rate of tag loss, we used a GLM of the log-transformed proportion of living adult females retaining their original p-Chip in a 7-day period against the number of days since the p-Chip was originally glued (setting this proportion to 1 for day 0, when the tagging occurred), weighted by simultaneous population size. Due to suspected differences in glue batch quality, this analysis was repeated with females split based on whether they were initially tagged in 2021 (*n* = 599) or in 2022 (*n* = 349), where 2021 corresponds to the first two vials of Surgi-Lock 2oc (including that used for the glue comparison study), and 2022 corresponds to a third vial. These analyses were limited to periods when at least 5% of the population was still alive.

### Effect of tagging on reproduction

To test whether the gluing of tags may have blocked the gonopores of females, we used a subset of female salmon lice that were tagged before (*n* = 55) or shortly after (*n* = 178) having extruded a first clutch of eggs. The number of egg matrices these females had extruded at the time of their initial tagging (0, 1 or 2) was compared to the maximum number they were observed to extrude within two observations (two weeks) after being tagged. Females that did not live through at least two observations or that were re-tagged within 15 days of their original tagging were excluded.

### Effect of tagging on mortality

Given that all adult female lice were tagged at the start of the study, no direct comparison of tagged *versus* untagged lice is made here (but see **Table SM1 & Figure SM3**). To estimate the impact of gluing adult females with 2oc on their mortality, we therefore compared females that were tagged once to those that were tagged more than once, with the reasoning that this constitutes a “second dose”. We selected all females in our dataset that were re-tagged seven days after receiving their original RFID tag (i.e. that were glued twice in two weeks), and limited to lice in 15-fish tanks (*n* = 49). For comparison, we selected a cohort of single-exposure females that were originally tagged on the same date and in the same tank as each of the selected re-tagged females and therefore at a similar stage of maturation, and which also survived to at least seven days after receiving their original RFID tag and were never re-tagged (*n* = 270).

To test whether multiple exposures to 2oc had long-term impacts on adult female mortality, we compared the number of days during which female lice lived after being re-tagged to that of females only tagged once using a Cox Proportional Hazards model. To test for shorter-term effects of the glue on female mortality, we used a chi-square test to compare the numbers of dead and live females after 14 days, according to whether they had been re-tagged or not.

All statistical analyses and visualizations of the data were performed in the statistical programming environment R (version 4.2.3, R Core Team 2023), using the R packages *broom* (Robinson et al. 2023), *survival* (Therneau 2023), *ggeffects* (Lüdecke 2018), and *tidyverse* (Wickham et al. 2019).

Models were validated by checking for overdispersion of residuals. For the Cox Proportional Hazards model, the package *survival* (Therneau 2023) was used, and diagnostic plots were generated with the package *survminer* (Kassambara et al. 2021). Model results reporting for glm functions used the package *report* (Makowski et al. 2023).

## Results

### Comparison of two alternative types of glue

Almost all females could be assigned as either having retained or lost their tag, with only two females in the 2oc group and four in the e2c group unaccounted for. Tag retention decreased significantly over time (glm; days since tagging: *b* = -0.006, 95% CI: -0.01 – -0.003, *p* < 0.001). In addition, retention decreased significantly faster for lice glued with e2c than for those glued with 2oc (glm; glue type * days: *b* = -0.01, 95% CI: -0.02 – -0.007, *p* < 0.001), although the main effect for glue type was not statistically significant (glm; glue type: *b* = 0.05, 95% CI: -0.1 – 0.19, *p* = 0.53; **Figure 2a, Table 1**).

**Table 1.** Generalised linear model parameter estimates of RFID tag retention for female salmon lice as a function of glue type (e2c vs. 2oc) and number of days since initial tagging. The model was fitted using the log-transformed proportion of lice recorded as retaining their tag, weighted by concurrent population size. See also Figure 2a.

|  | Estimate | SE | *t* | *P* |
| --- | --- | --- | --- | --- |
| Intercept | 0.008 | 0.051 | 0.156 | 0.877 |
| Glue type | 0.046 | 0.073 | 0.628 | 0.535 |
| Days since tagging | -0.006 | 0.002 | -3.364 | 0.002 |
| Glue type x Days | -0.012 | 0.003 | -4.657 | 5.72e-5 |

**Figure 2.**
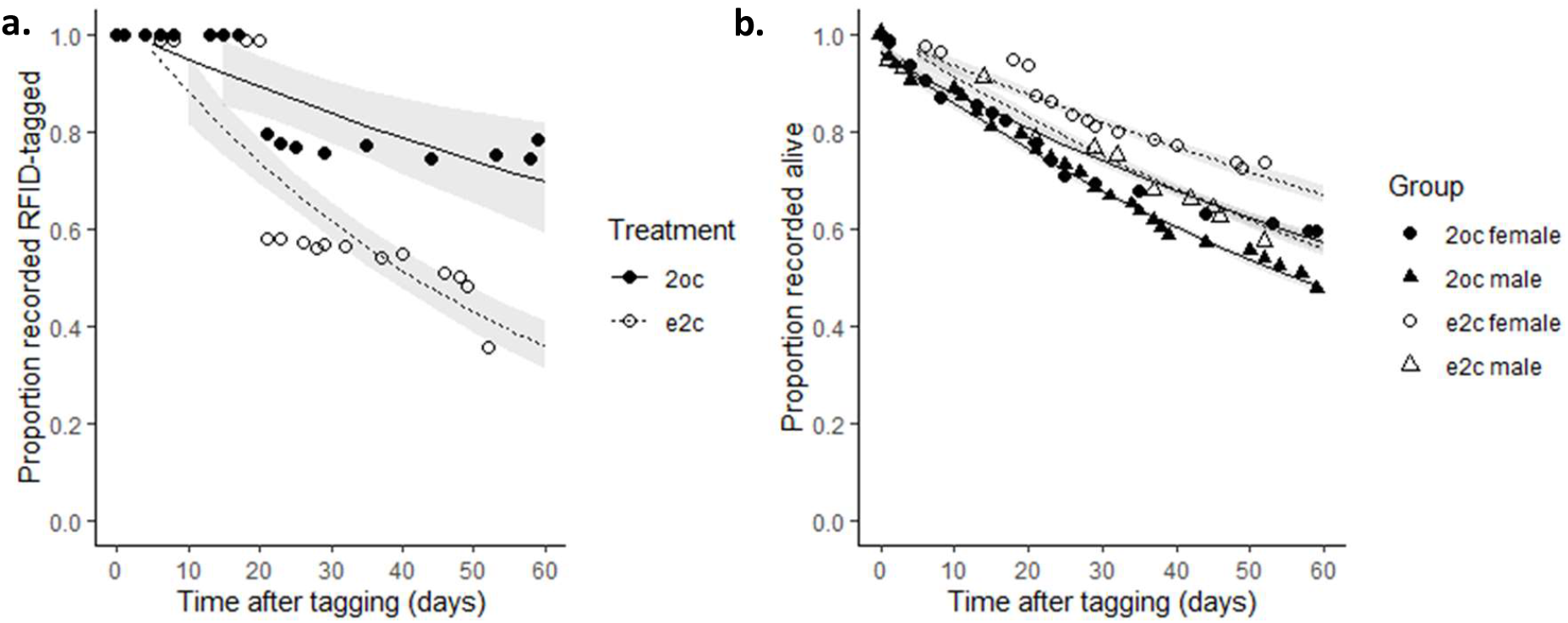
Comparison of (a) RFID tag retention including observations from tank outlet filters and (b) the proportion of living adult male and female lice according to the type of glue used for tagging the female salmon lice (e2c = ethyl 2-cyanoacrylate, 2oc = 2-octyl cyanoacrylate). Day 0 is the day when RFID tags were glued dorsally on the abdominal segment of adult female salmon lice. Males were not tagged but are included under the glue type used on females in the same tank.

The proportion of living salmon lice in all groups decreased over time (glm; days since tagging: *b* = - 0.009, 95% CI: -0.009 – -0.008, *p* < 0.001). Although the overall log-proportion of lice alive did not differ between the sexes (glm; sex: *b* = 0.007, 95% CI: -0.02 – 0.03, *p* = 0.61), males were lost at a significantly higher rate than females (glm; sex * days: *b* = -0.003, 95% CI: -0.004 – -0.002, *p* < 0.001). The proportion of lice alive (both males and females) was higher in the tank where the females were tagged using e2c than in the tank where females were tagged using 2oc (glm; glue type: *b* = 0.04, 95% CI: 0.02 – 0.07, *p* = 0.001), but decreased at a significantly lower rate (glm; glue type * days: *b* = 0.002, 95% CI: 0.001 – 0.003, *p* < 0.001; **Figure 2b, Table 2**).

**Table 2.**
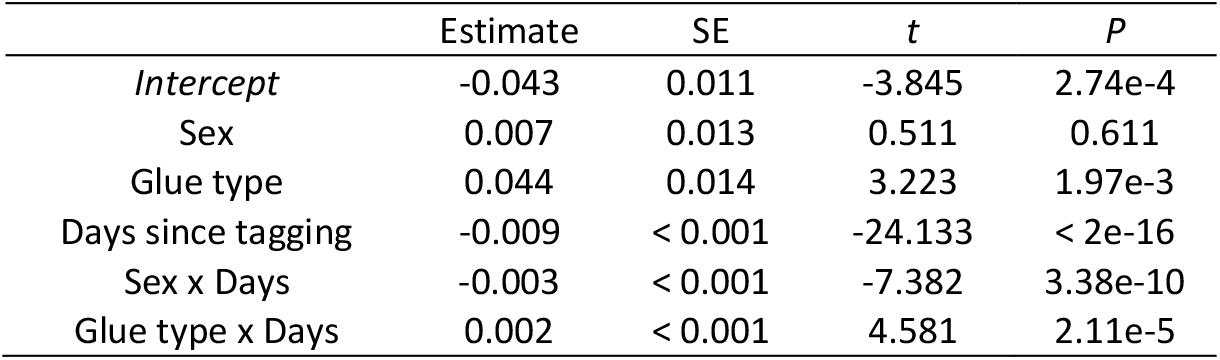
Generalised linear model parameter estimates of the mortality over time for tagged female salmon lice and untagged male salmon lice as a function of sex (males vs. females), the glue type used on the females in that tank (e2c vs. 2oc), and the number of days since initial tagging. The model was fitted using the log-transformed proportion of lice recorded as living, weighted by starting population size. See also Figure 2b.

### Documenting retention time

During this study (from 11th January 2021 to 16th December 2022), 216 of 948 p-Chips glued using 2oc were replaced (i.e., 22.7%); of these, four were replaced due to loss of function, while the remaining 212 had detached from the louse. The proportion of tagged individuals declined exponentially over time (glm of log-transformed proportion; days since tagging: *b* = -0.003, 95% CI: --0.004 – -0.003, *p* < 10^−4^; **Figure 3, Table 3**). Time until tag loss ranged from 0 to 150 days, while the longest time that a female louse from this study retained its original p-Chip was 263 days (until death).

**Table 3.**
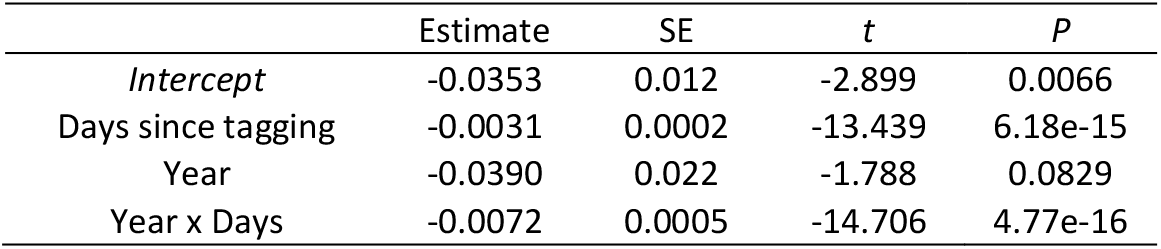
Generalised linear model parameter estimates for the log-transformed proportion of living female salmon lice retaining their original RFID tag as a function of the year during which they were tagged (2022 vs. 2021) and the number of days since initial tagging, with the proportion at day 0 set to 1 and weighted by population size. See also Figure 3.

**Figure 3.**
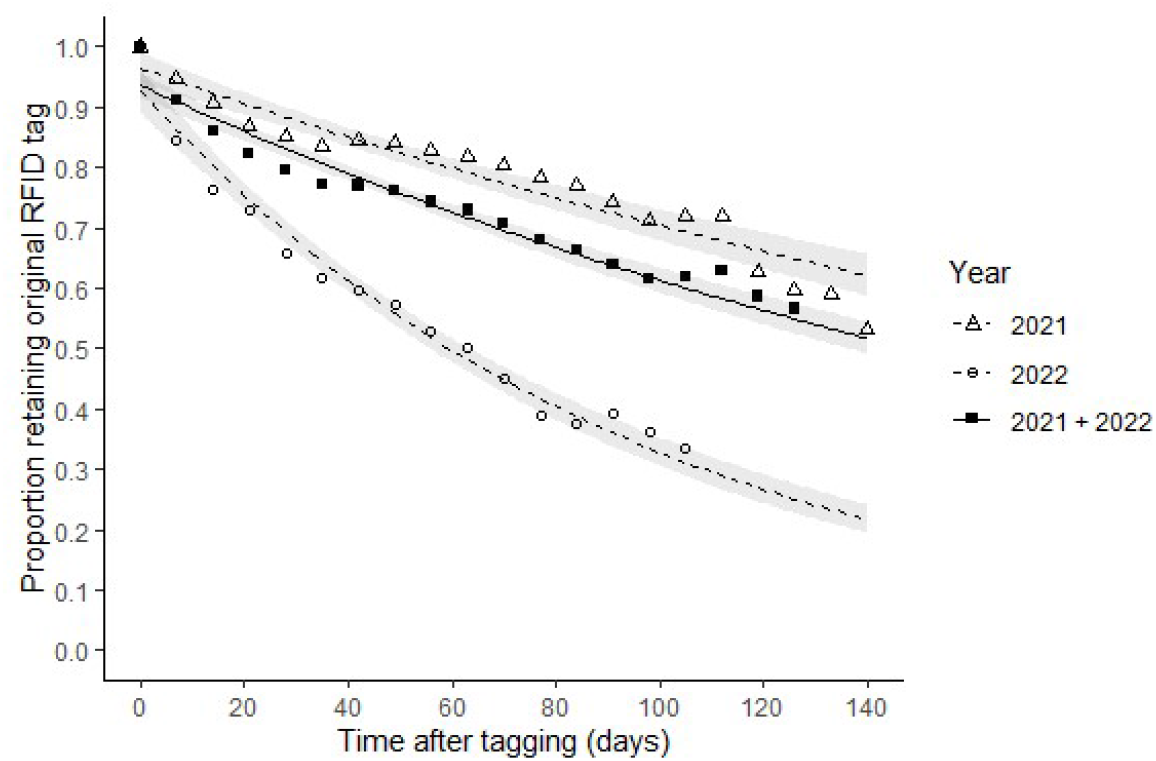
The proportion of living female salmon lice retaining their original RFID tag decreased over consecutive 7-day intervals, starting from the date of initial tagging (day 0, *n* = 948) up to 126 days (*n* = 53). The exponential regression line for all females is shown in solid black. There was evidence of batch differences in the glue used, since the rate of decrease in RFID tag retention for one vial used in 2022 (*n* = 349) was significantly higher than for two vials used in 2021 (*n* = 599). The exponential regression lines for these subgroups are shown as dashed lines.

Comparing the two vials of glue used in 2021 to the one used in 2022, all the same brand but purchased from different suppliers, we found that in 2021, 117 of 599 p-Chips were lost (i.e., 19.5%), while 99 of 349 p-Chips were lost in 2022 (i.e., 28.4%). While the log-transformed proportion of lice retaining their original tags did not differ significantly between years (glm ; year: *b* = -0.04, 95% CI: -0.08 – 0.004, *p* = 0.07), it decreased more rapidly in 2022 than in 2021 (glm ; Year * Days since tagging: *b* = -0.007, 95% CI: -0.008 – -0.006, *p* < 0.001; **Figure 3, Table 3**).

### Effect of tagging on reproduction

Of the 233 female salmon lice checked for differences in egg extrusion before and after attachment of a p-Chip, all lice were observed to be able to extrude eggs from at least one gonopore except for two females that were never observed to produce eggs. Of the lice glued before they extruded eggs (*n* = 55), 51 extruded a pair of egg matrices within two weeks, while two extruded only one egg matrix, and two extruded zero. Of the lice glued after they had extruded two egg matrices (*n* = 172), 164 continued to extrude pairs, while six extruded only one egg matrix and two did not extrude any eggs in that 2-week period. For a subset of females glued after they had extruded only one egg matrix (*n* = 6), two continued to extrude only one, while four extruded pairs.

### Effect of tagging on mortality

We found no significant difference in long-term survival for females that were re-tagged seven days after their original tagging compared to those that were only tagged once (Cox Proportional Hazard, *n* = 319: HR = 0.94, 95% CI: 0.69 – 1.28, *p* = 0.693). After 14 days, 32 of 49 re-tagged and 175 of 270 non-re-tagged females were still alive. A chi-square test found no significant difference in short-term mortality between these groups (*χ*^2^ = 6.65e-30, df = 1, *p* = 1).

## Discussion

We compared two types of glue for attaching p-Chip RFID tags to adult female salmon lice. We found that 2-octyl cyanoacrylate (Surgi-Lock, 2oc) resulted in a significantly higher retention rate than ethyl 2-cyanoacrylate (Würth Klebfix, e2c). Mortality was found to be significantly higher for adult lice (both males and females) in the tank where females were glued with 2-octyl cyanoacrylate than in the tank where females were glued with ethyl 2-cyanoacrylate. This result was surprising as males are not tagged and thus not in direct contact with the glue. Even though this might be due to 2-octyl cyanoacrylate having a certain level of toxicity that might also have affected the males living in the same tank (same water), we cannot rule out a mere tank effect. Due to logistical constraints and since we had not yet established that adult lice could also often be identified from pictures (**Figure SM1**) when this study started, we were not able to have additional tanks or mix treatments within tanks. The two types of glue were therefore compared in distinct tanks where the salmon hosts might have experienced slightly different levels of stress or behaved otherwise differently, causing the lice to detach from their hosts at different rates. This interpretation is supported by previous reports of unexplained between-tank variability in the time during which adult female lice stayed on their host at high host densities (Hamre et al., 2009). In addition, a more recent analysis of data collected prior to this study did not indicate toxicity on tagged adult female lice for either glue when compared to untagged females (**Table SM1, Figure SM3**).

During a more extensive period of testing, we further investigated the impact of 2-octyl cyanoacrylate on female salmon louse mortality by selecting a subset of females that had been re-tagged seven days after their initial tagging and comparing these to a cohort of females, from the same tanks and at a similar stage of maturity, that had never been re-tagged. We reasoned that if 2-octyl cyanoacrylate were toxic, then this would represent a “double dose” within a relatively short period of time. There was no significant difference in the remaining lifespan of these two groups. We also tested for increased mortality in the two weeks immediately following two consecutive exposures but did not find evidence of increased mortality in this period either. Additionally, 2-octyl cyanoacrylate was more convenient to use, as it remained workable for a longer period in open air than ethyl 2-cyanoacrylate, preventing accidental adhesions of tools. Given the purpose and scale of our planned research, the longer tag retention and better tractability of 2-octyl cyanoacrylate were substantial enough to justify using it in subsequent experiments.

We found that gluing had little apparent impact on the extrusion of eggs for female salmon lice. The majority of lice (219 of 233) that were glued after producing zero, one, or two egg matrices (strings) went on to produce a full pair, regardless of the initial number. Closer examination of pictures of the two lice that never extruded egg matrices revealed that one of them had eggs in its ovaries that could not be extruded and died shortly after, while for the other one, the ovaries remained empty. Of the eight females that went from extruding two egg matrices to extruding zero or one, four resumed producing pairs while the other four died shortly after, one of those with glue residue visible around the gonopore. A small subset of lice appeared to either naturally extrude egg matrices from only one gonopore or to produce matrices that naturally detached shortly after extrusion. Glue residue was visible for three females tagged prior to extruding eggs and that later extruded from only one side, amongst which one resumed producing a pair of matrices shortly after. For those females that could be recaptured within two weeks, our study did not detect a significant effect of tagging on their further survival or reproduction. All in all, our study did not find strong evidence for any impact of the tagging procedure on vital or reproductive functions. Testing the effects of tagging on other aspects of the biology of the species may however be useful for studies differing from ours in topic or goal.

Over a longer period and including all batches of glue, we found that p-Chips glued to the abdominal segment of adult female salmon lice with Surgi-Lock 2oc were lost at an apparently stable rate over time, as reflected by the negative exponential decline in the proportion of lice that retained their tags. This decline was rather shallow (although somehow greater during the first week), with a generally high percentage of lice retaining their original tag (about 82.5% after 30 days, or 72.6% after 60). We also found, however, that this rate may be affected by the age of the glue, or that there may also be differences in batch quality. After noticing an increase in lost p-Chips after opening a new vial of Surgi-Lock 2oc at the start of 2022, which was then also used for longer after opening, and comparing the retention rate for females initially tagged in 2021 to those tagged in 2022, we found a higher rate of decrease for tag retention in 2022. Subsequently, while we have not extended the analysis further, switching to a fourth vial in 2023 with a later manufacturing date resulted in a marked decrease in the number of tags replaced each week. This suggests that the vial used in 2022 was atypical, and that the normal retention rate for this glue may be closer to the regression line for 2021 (i.e. 87.8% retention after 30 days and 79.9% retention after 60; **Figure 3, Table 3**). In addition, although not documented here, retention might decrease as time elapses since opening a vial, so for the sake of caution we would recommend using vials of glue that are less than six months old. It may be feasible to develop a test for glue efficacy prior to use in research, such as by monitoring adherence to a suitable substrate submerged in water conditions approximating the study environment.

The life span of adult salmon lice in the wild has not been quantified, but in laboratory studies females are found to live for up to 191, 452, and 303 days post-infection (Heuch et al., 2000, Hamre et al., 2009, and Folk & Mennerat 2023, respectively). The rate of loss for female salmon lice in the lab appears positively related to stocking density, as well as the rates at which both males and females move among hosts (Mennerat et al., *in prep*.; see also Hamre et al., 2009). In the present dataset, the median female lifespan ranged from 22.5 to 50 days post-tagging in 15-fish and 5-fish tanks respectively. RFID tags may therefore cover the largest portion of an adult female’s life, especially in densely-stocked host populations, *i*.*e*. when it is the most time-consuming to identify individuals by other means.

Although the use of p-Chips requires some level of training, it appears to be a robust yet simple technology that allows for individual studies of ectoparasites, including those living in seawater like *L. salmonis*. More generally, this procedure for RFID tagging with microtransponders could easily be adapted for other aquatic and terrestrial species and be used as a relatively rapid, non-invasive, cost-effective method (compared to molecular approaches) for identifying invertebrates traditionally not considered suitable for individual studies, provided that they are sufficiently large and possess an exoskeleton.

Potential applications of this methodology for ecological and evolutionary research include facilitating data collection for behavioural and dispersal studies, thus potentially giving insight into a broader range of insects and other invertebrates. RFID tagging also allows documenting individual life histories, through which greater understanding of contemporary evolution can be gained using species not traditionally considered suitable for monitoring studies. This can prove particularly insightful in the fields of disease ecology and host-parasite coevolution where more empirical research is warranted, especially given the global environmental changes parasites are facing and the increasing disease threats linked to farming practices and other major human activities (Mennerat et al., 2010, Lafferty et al., 2015, Kennedy et al., 2016, Mennerat et al., 2017, Ugelvik et al., 2017).

## Ethical statement

All procedures adhered to “the Regulation on the Use of Animals in Research” under the Norwegian Food Safety Authority, and the Atlantic salmon in this study were used according to a permit from the Norwegian animal research authority Mattilsynet to A. Mennerat (FOTS id 24917).

## Acknowledgements

We are grateful for help with lice acquisition to Kjetil Stensland, Haldor Haugsdal, Tina Oldham, Samantha Bui, and Frode Oppedal. Thanks to Per Gunnar Espedal and Lars Are Hamre for daily fish feeding and maintenance (including during lockdown), to Richard Telford for statistical input and to Mikko Heino for discussions. Additional thanks to Freya Coursey, Renate Løvlund Andersen, and Peder Moberg for assistance with weekly data collection.

## Data, scripts, code, and supplementary information availability

Data, scripts, and code are available online: https://doi.org/10.5281/zenodo.10406878 (Folk & Mennerat 2023).

## Appendix Supplementary materials

**Figure SM1.**
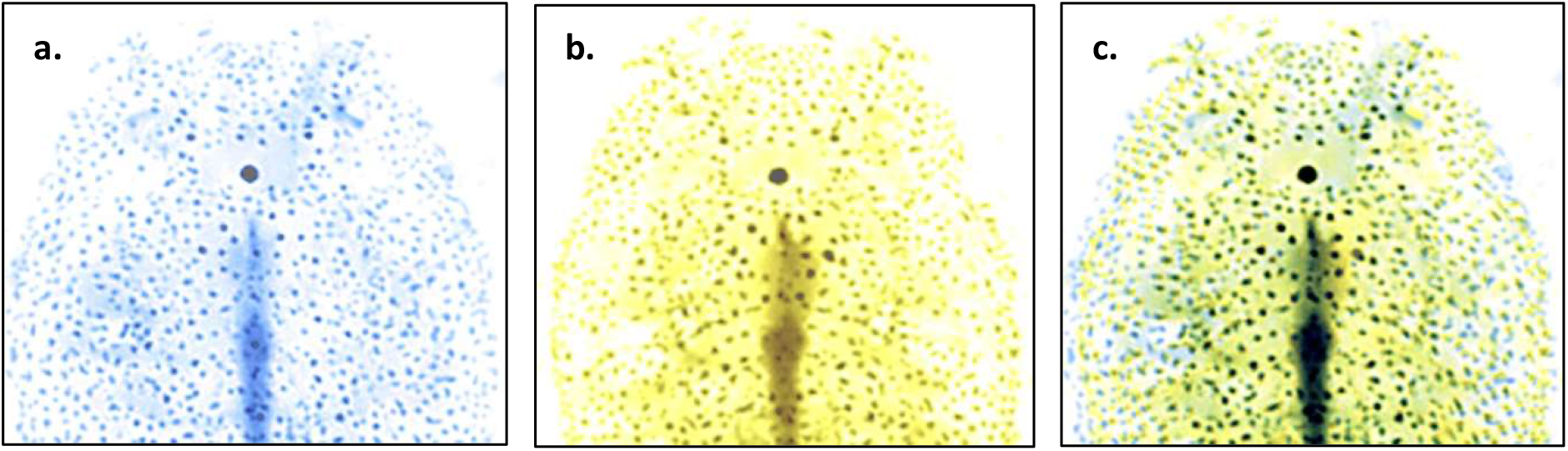
An illustration of conserved pigmentation pattern in adult female salmon lice. The same female is shown here (a) in blue at 61 days post-infection and (b) in yellow at 271 days post-infection. The central dark stripe is the gut. In (c) the two images are overlaid; areas in black thus indicate a high degree of overlap.

**Figure SM2.**
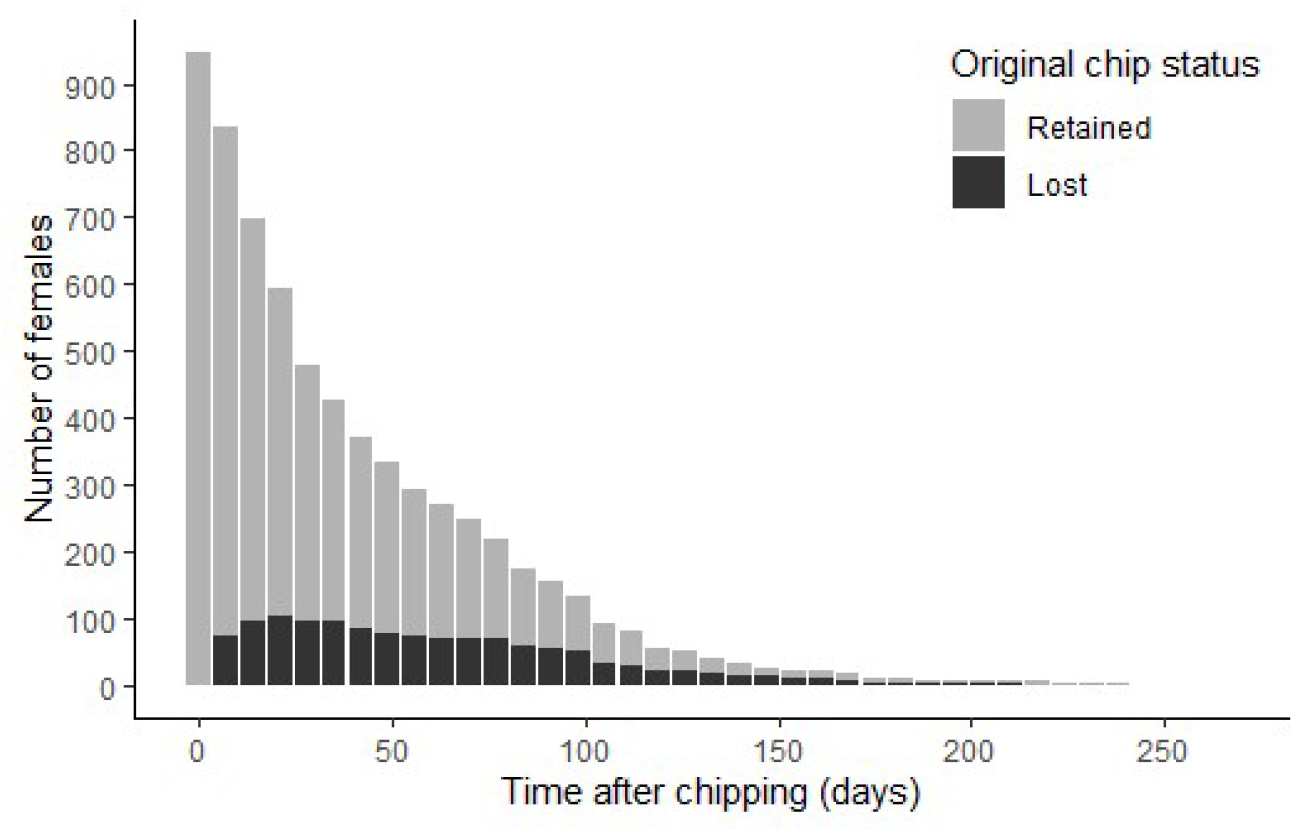
The number of living female salmon lice in 7-day intervals from the date of initial tagging (day 0, *n* = 948) up to 266 days (*n* = 1); the colours indicate whether females had lost their original RFID tag (black) or retained it (grey).

**Figure SM3.**
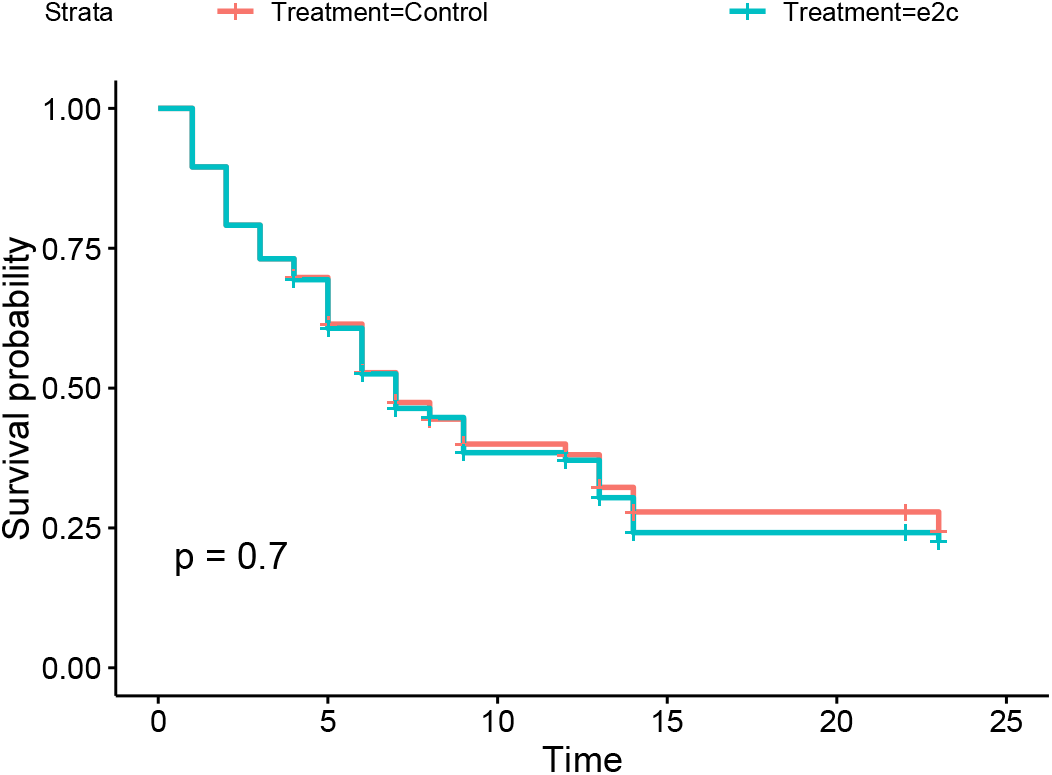
Results from an earlier pilot study showing no significant difference in survival for female lice tagged using *e2c*, as compared to untagged controls. Adult female salmon lice originating from the lab strain *Ls1* were collected on 22/11/2017 from their hosts (with help from Lars Hamre, University of Bergen). On the same day they were either tagged (n=28) or handled similarly except for tagging (n=28). They were then immediately placed in individual wells under similar conditions of temperature and water flow, after which their survival was monitored for 23 days (note that in this case they were detached from their hosts and thus could not feed). Survival was compared between treatments using Kaplan-Meier survival analysis (using the *survival* R package).

**Table SM1.** Results from a preliminary study showing no difference in survival for female lice tagged using *2oc*, as compared to untagged controls. 16 salmon hosts in a single 500L tank could be infected on 18/3/2020 despite lockdown, with salmon lice larvae hatched from eggs collected 10 days before at Austevoll, Norway (about 30 larvae / fish). At day 63 post-infection adult females were collected from their hosts and either tagged using 2oc (n=44) or left untagged (n=45). The tagging treatment was assigned randomly (about every second female was tagged on each host). After being photographed, the lice were placed back on their respective hosts, and the hosts back in their common tank. At day 34 post-tagging all females were collected again, photographed and immediately placed back on their hosts. All hosts were euthanised at day 100 post-tagging and females were photographed one last time. The tank outlet system was equipped with a similar filter as described in this article, and checked at 2-3 days intervals at most. Female lice sampled on days 34 and 100, as well as those found in filters, were identified from pictures whenever possible by comparing their pictures to those taken on tagging day (day 0). The numbers of lice identified as initially tagged *vs*. untagged are reported below. There is no indication that the numbers of tagged lice decreased more rapidly than those for untagged lice.

|  | Initially untagged | Initially tagged |
| --- | --- | --- |
| Day 0: tagging | 45 | 44 |
| Day 34: intermediate sampling | 22 | 28 |
| Day 100: final sampling | 14 | 17 |
| Dead in filters & successfully identified | 8 | 3 |

